# Three-gene PCR and high-resolution melting analysis for differentiating vertebrate species mitochondrial DNA for forensic and biodiversity research pipelines

**DOI:** 10.1101/636605

**Authors:** Daniel O. Ouso, Moses Y. Otiende, Maamun Jeneby, Joseph W. Oundo, Joel L. Bargul, Scott Miller, Lillian Wambua, Jandouwe Villinger

**Affiliations:** International Centre of Insect Physiology and Ecology (*icipe*), P.O Box 30772 – 00100, Nairobi, Kenya; Biochemistry Department, Jomo Kenyatta University of Agriculture and Technology (JKUAT), P.O Box 62000 – 00200, Nairobi, Kenya; Kenya Wildlife Service, Veterinary Department, P.O Box 40241-00100, Nairobi, Kenya; Institute of Primate Research, National Museums of Kenya, Department of Tropical and Infectious Diseases, P. O. Box 24481-00502, Karen, Nairobi, Kenya; National Museum of Natural History, Smithsonian Institution, Washington, DC, USA

## Abstract

Reliable molecular identification of vertebrate species from morphologically unidentifiable tissue is critical for the prosecution of illegally-traded wildlife products, conservation-based biodiversity research, and identification of blood-meal hosts of hematophagous invertebrates. However, forensic identification of vertebrate tissue relies on the sequencing of mitochondrial cytochrome oxidase I (*COI*) ‘barcode’ genes, which remains costly for purposes of screening large numbers of unknown samples during routine surveillance. Here, we adopted a rapid, low-cost approach to differentiate 10 domestic and 24 wildlife species that are common in the East African illegal wildlife products trade based on their unique high-resolution melting profiles from *COI*, cytochrome b, and *16S* ribosomal RNA gene PCR products. Using the approach, we identified (i) giraffe among covertly sampled meat from Kenyan butcheries, and (ii) forest elephant mitochondrial sequences among savannah elephant reference samples. This approach is being adopted for high-throughput pre-screening of potential bushmeat samples in East African forensic science pipelines.

## Introduction

Unsustainable hunting, consumption, and sale of bushmeat in Africa contribute immensely to the decline of threatened wild animal species. The global bushmeat trade is valued at several billion dollars. Up to 270 ton of bushmeat were flown into Europe through a single airport in 2010 from Africa^1^. While this is a major crisis for wildlife in central and western Africa, it is a growing concern in eastern and southern Africa^2^. Efforts to regulate or prevent illegal wildlife trade depends on accurate, efficient, and sustainable tools for species identification of confiscated and surveillance samples.

Illegal wildlife trade is mainly fueled by the need for diet and income supplementation^3^. The consequences of direct human contact with bushmeat have been severe. A classic example of disease originating from or harbored by bushmeat includes is Ebola virus disease, which has infected humans upon contact with infected wild animals such as fruit bats, nonhuman primates, and forest antelopes^4,5^. The impact of bushmeat hunting on animal populations can also be severe^6^. Many favored wild animal species for bushmeat are already endangered, some close to extinction^7^. There are also flow on effects to the ecosystems^8^ and tourism.

Concerted efforts have been put in place to save precious flora and fauna, including awareness campaigns and fencing of parks and conservancies. However, cases of bushmeat hunting are still rampant even with laws prohibiting bushmeat trade. Law enforcement can only be effective when backed by efficient prosecution, which relies on proper surveillance, concrete evidence, and informed policies. Accurate identification of suspect samples forms the basis of forensic evidence, which currently relies widely on sequencing of barcode cytochrome c oxidase sub-unit I (*COI*) genes^9,10^. This approach is increasingly becoming accepted and adopted as a means of court-admissible evidence generation for wildlife crime prosecutions in East Africa. However, as only a small proportion of potential samples sequenced are of illegally traded wildlife products, the cost of surveillance by mass barcode sequencing is high and thus not sustainable in the long term.

DNA barcode sequencing has replaced earlier molecular methods of vertebrate species identification, such as restriction fragment length polymorphism^11^, random amplification of polymorphic DNA^12^, and amplified fragment length polymorphism^13^, which suffer poor reproducibility and thereby limit the development of reference databases^14^. More recently, sequence tag repeats^15^ and single nucleotide polymorphism^16^ have been used, which are however limited to detecting very specific wildlife species.

However, high-resolution melting (HRM) analysis has been used with a number of genes to identify species among diverse viruses^17^, bacteria^18^, malaria *Plasmodium*^19^, mosquitoes^20^ and their bloodmeals^21^, plant products^22^, and animals within discrete families^23^, as well as human individuals^24^. HRM analysis is a fast, sensitive, and specific tool developed for genotyping PCR product sequence variations^25^ that employs the use of intercalating fluorescent dyes, such as EvaGreen^26^. The dyes undergo rapid solvent fluorescence quenching as the duplex DNA PCR products are melted. The amplicon melting temperatures (T_m_) and specific melt curve shapes are dependent on DNA complementarity, G-C content, and amplicon length. HRM analysis has not been standardized to support forensic pipelines for identifying illegally traded wildlife products.

Therefore, to address this lack of cost-effective techniques to robustly identify vertebrate species from morphologically indistinct samples, we adopted and validated an HRM analysis-based approach to rapidly identify and differentiate domestic species from wild vertebrate species commonly targeted for bushmeat in East Africa^27^. By systematically comparing HRM profiles generated by *COI*, cytochrome b oxidase (*cyt b*), and *16S* ribosomal (r)RNA gene PCR products, we show that this approach can robustly differentiate domestic vertebrate species from wildlife species, allowing for forensic barcode sequence confirmations to be limited to only wildlife specimens. During the validation process, we made observations on the mitochondrial history of East African elephants and zebra populations. We further used the approach in a proof-of-concept study to identify illegal bushmeat among samples covertly purchased from butcheries in the Naivasha region of Kenya.

## Results

### Species-specific mitochondrial gene high-resolution melting analysis profiles

We generated HRM profiles from short mitochondrial *COI*, *cyt b*, and *16S rRNA* PCR products to differentially identify the species of 107 reference tissue samples (Supp Table 1), which included 10 domestic and 24 East African wildlife vertebrate species (18 Bovidae, four Equidae, four Felidae, two Elephantidae, two Rhinocerotidae, three Suidae, and one Camelidae species) (Figs. 1-5). Where available, we used multiple samples for particular species. Species identifications were further confirmed by sequencing of the *COI* barcode region. The domestic species, including three members of the Bovidae family (cattle-*Bos taurus*, goat-*Capra hircus*, and sheep-*Ovis aries*), one member each for Suidae (pig-*Sus scrofa*), Equidae (donkey-*Equus asinus*), Camelidae (camel-*Camelus dromedarius*), and Leporidae (rabbit-*Oryctolagus* sp.), and two Phasianidae (turkey-*Meleagris gallopavo* and chicken-*Gallus gallus*), were successfully differentiated from all other wild animal specimens by three-marker PCR-HRM analysis.

**Figure 1:**
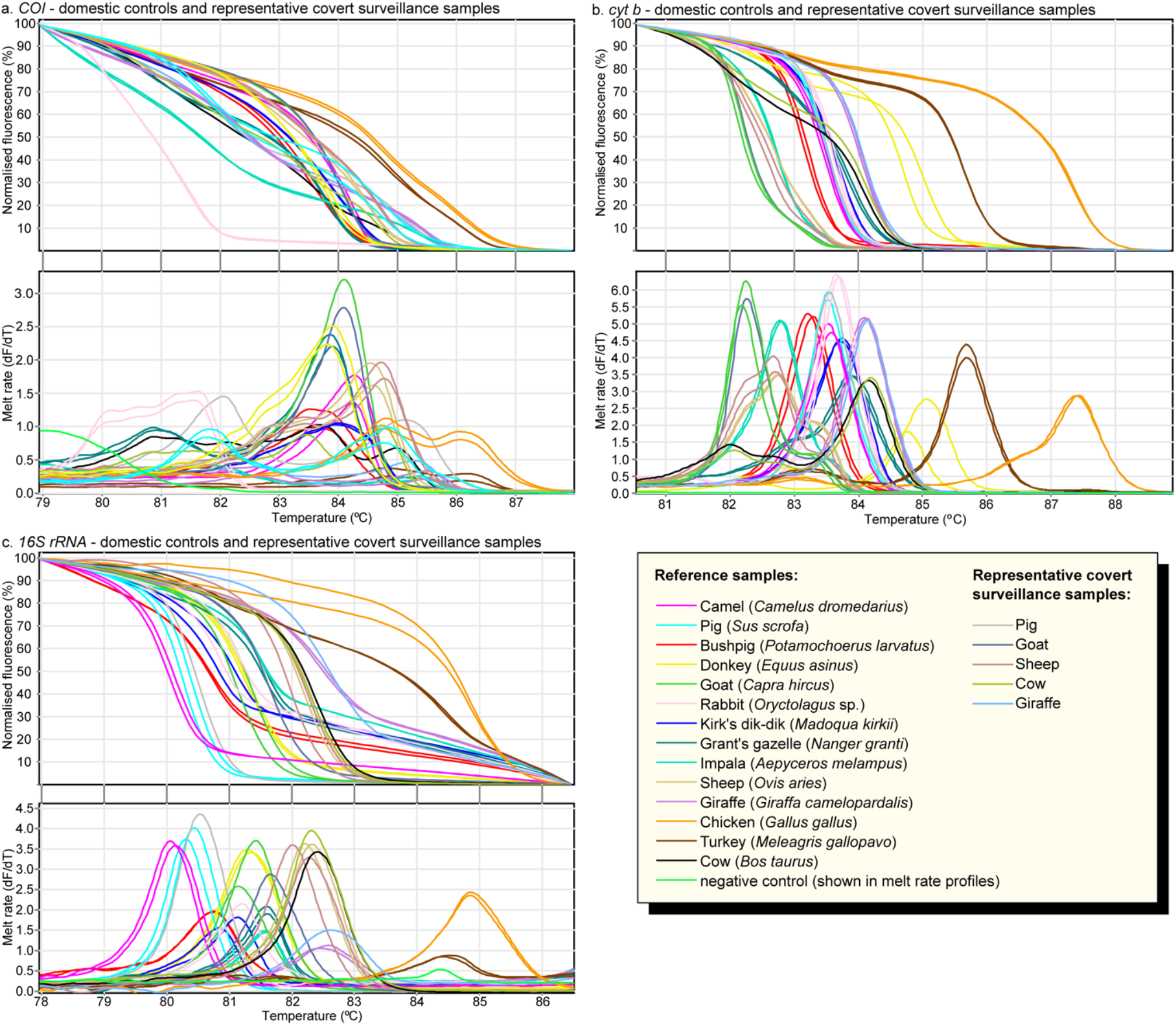
Distinct normalized HRM and melt rate profiles of domestic reference and representative covert surveillance samples. Normalized HRM profiles are represented as percent fluorescence and melt rates are represented as change in fluorescence units with increasing temperatures (dF/dT) for (a) *COI* (b) *cyt b* and (c) *16S* rRNA markers.

Though most species could be differentiated by pairwise comparisons of HRM profiles using all three markers, some species could only be differentiated by one or two of the three markers (Fig. 6) due to similar HRM profiles within 1°C melting temperature (T_m_) ranges or due to poor amplification of some species with the primers of particular markers. For example, waterbuck (*Kobus ellipsiprymnus*) failed to amplify with *COI*, but could be differentiated from all other species based on its *cyt b* and *16S rRNA* HRM profiles. Some species showed similarities in both shapes and melting temperature for particular markers. For example, among *COI* HRM profiles, pig samples generated similar *COI* and *cyt b* HRM profiles within a 1°C T_m_ range to those generated by giraffe (*Giraffa camelopardalis*) samples, but could be clearly differentiated based on their distinct *16S rRNA* HRM profiles (Figs. 2 and 6).

**Figure 2:**
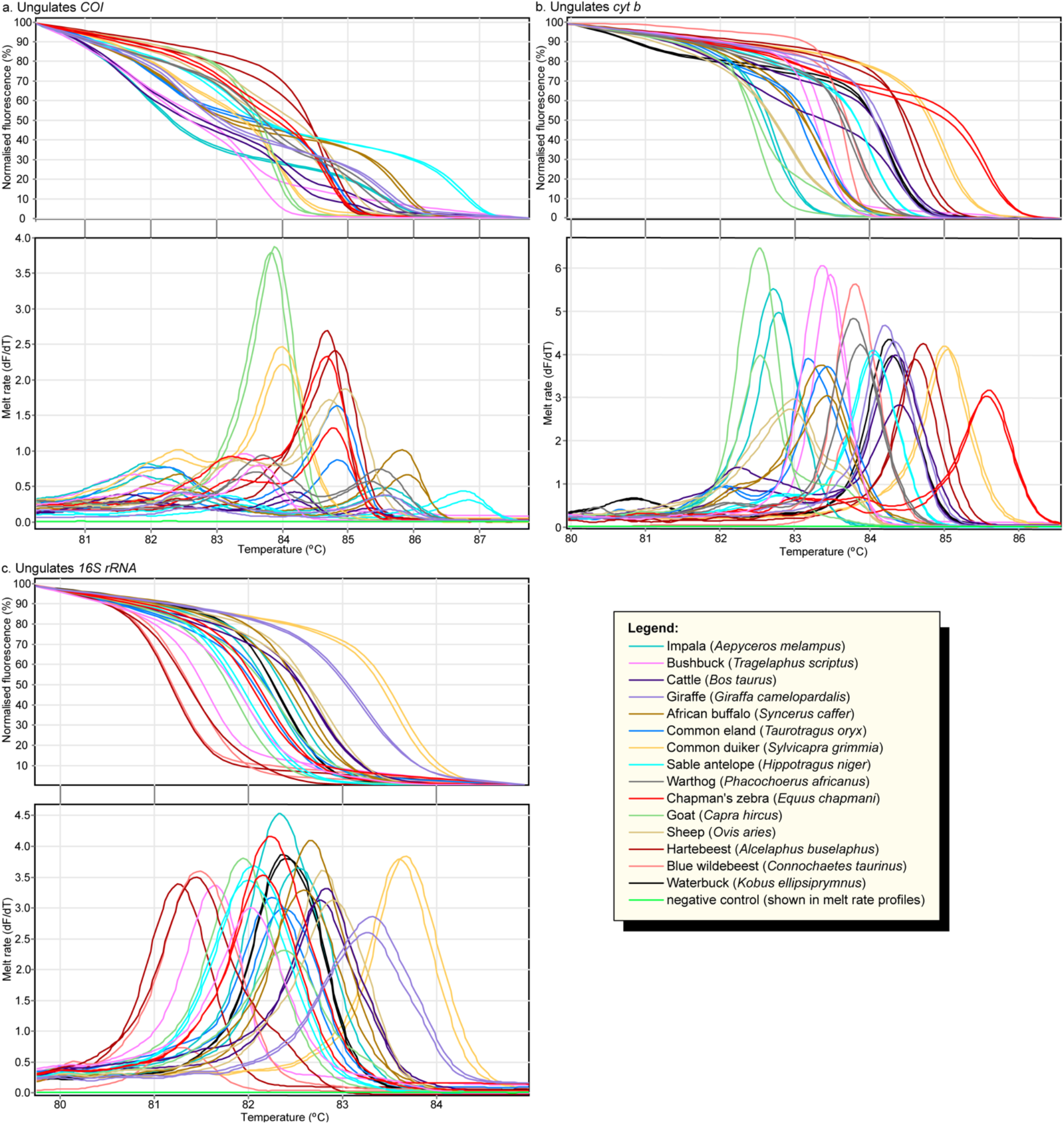
Distinct PCR-HRM profiles of ungulate species. Normalized HRM profiles are represented as percent fluorescence and melt rates are represented as change in fluorescence units with increasing temperatures (dF/dT) for (a) *COI* (b) *cyt b*, and (c) *16S* rRNA markers.

We generated two distinct sets of HRM profiles for elephant reference samples obtained from Kenya Wildlife Service (KWS) using all three markers (Fig, 3). Upon COI barcode sequencing of these samples, we found that one set of HRM profiles corresponded to the expected savannah elephant (*Loxodonta africana*), which is endemic to Kenya (GenBank accession xxx). Interestingly, the other set of HRM profiles were generated from savannah elephants with forest elephant (*Loxodonta cyclotis*) mitochondrial DNA (mtDNA) (GenBank accession xxx), not endemic in Kenya. All markers were also able to distinguish the two species of rhinos, black (*Diceros bicornis*) and white (*Ceratotherium simus*) rhinos (Fig. 3). Among equine samples, zebra species endemic to East Africa (plains zebra-*Equus quagga*, Chapman’s zebra-*Equus chapmani*, Grévy’s zebra-*Equus grevyi*) and donkey (Fig. 4), as well as available Felidae reference samples (cheetah-*Acinonyx jubatus*, leopard-*Panthera pardus*, lion-*Panthera leo*, domestic cat-*Felis catus*) (Fig. 5), could be clearly distinguished by three-marker PCR-HRM from each other and other domestic and wildlife species (Figs.1-3,6). During early HRM experiments using DNA extracts provided by KWS, we also obtained unique PCR-HRM profiles for loggerhead sea turtle (*Caretta caretta*) and green sea turtle (*Chelonia mydas*) (Supp. Fig. 1).

**Figure 3:**
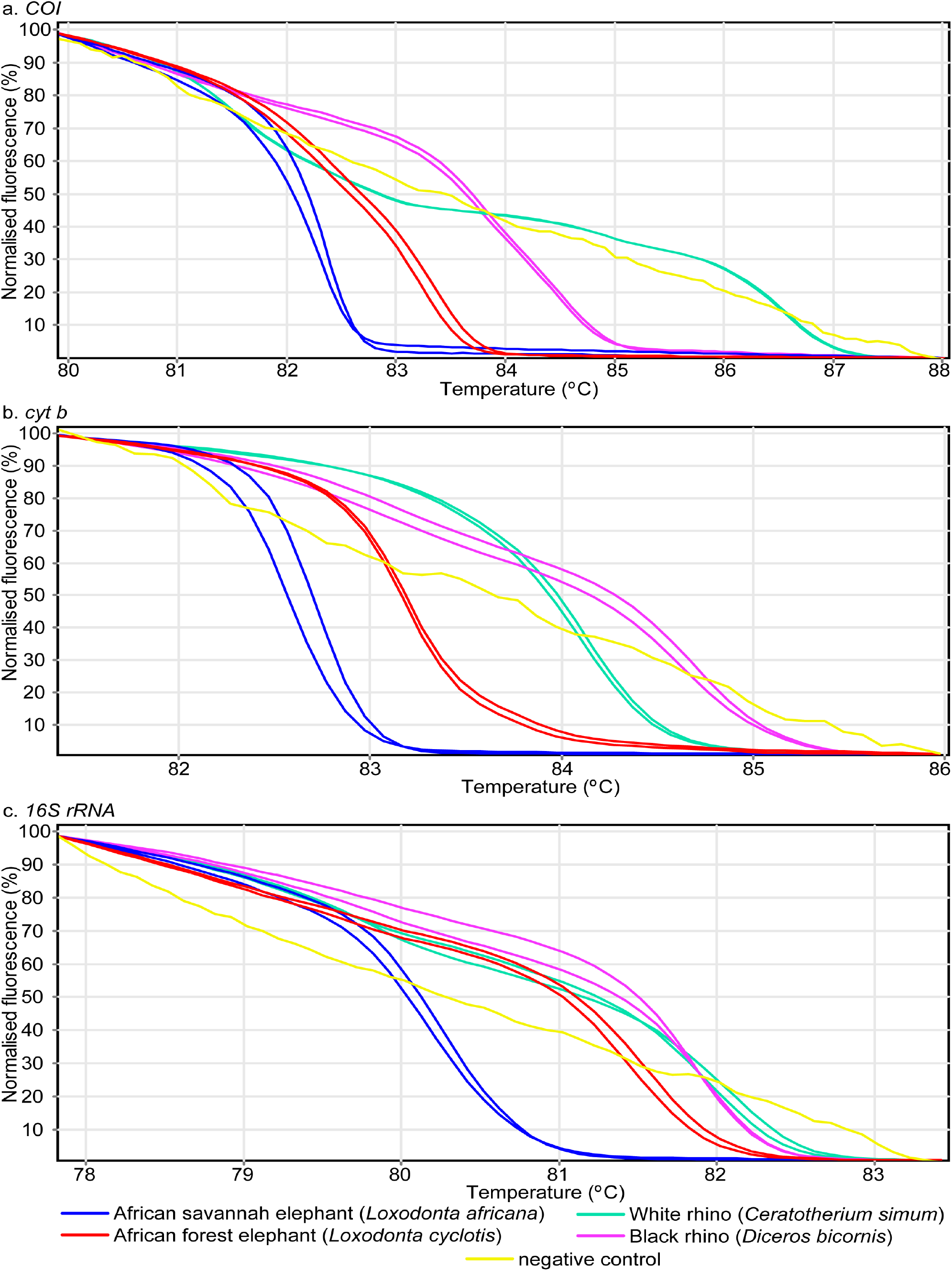
Distinct PCR-HRM profiles among Elephant and Rhino species. Normalized HRM profiles are represented as percent fluorescence for (a) *COI* (b) *cyt b*, and (c) *16S* rRNA markers. The African forest elephant mitochondrial amplicons were obtained from African savannah elephant reference samples.

**Figure 4:**
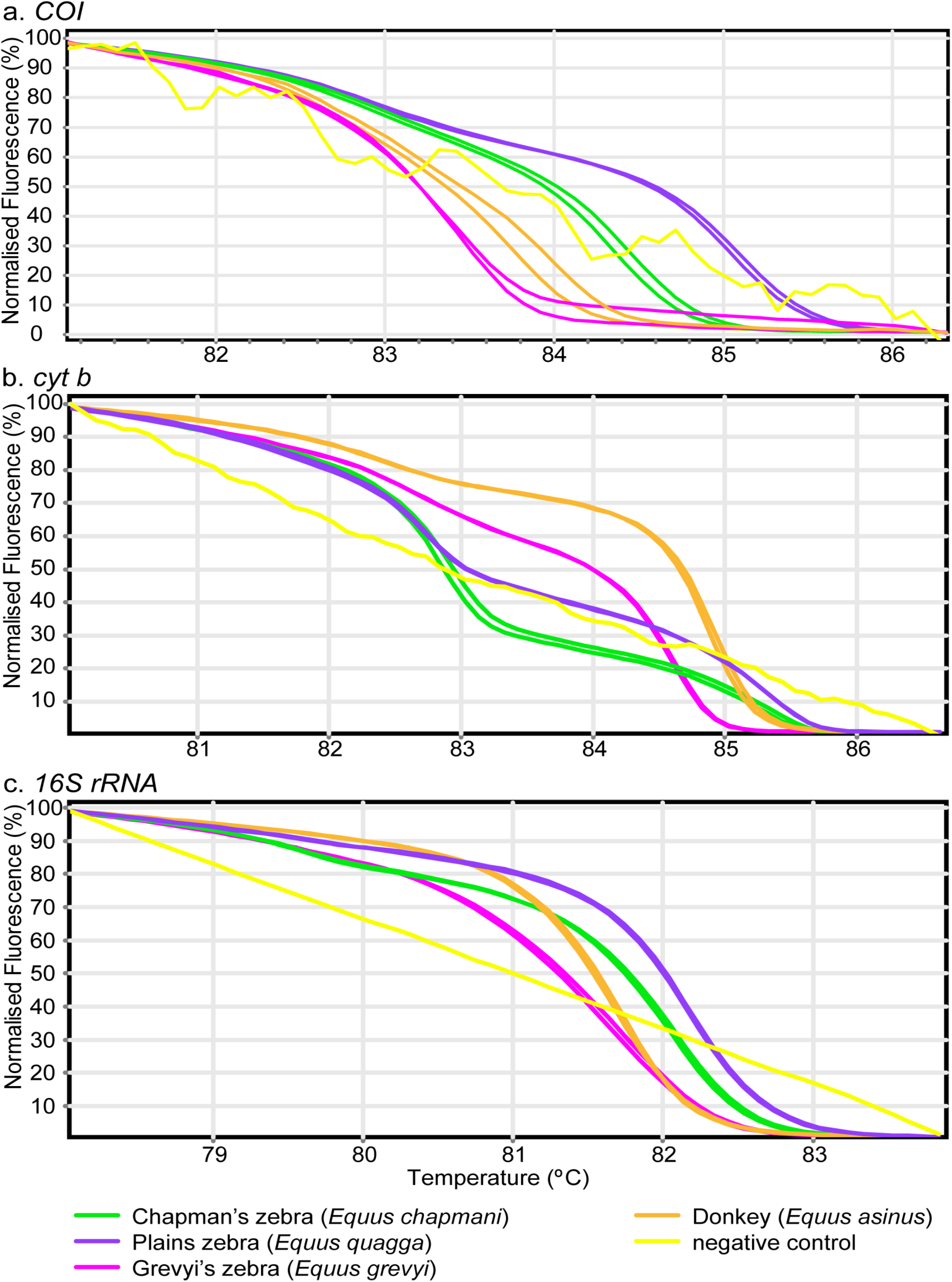
Distinct PCR-HRM profiles for the Equidae family showing the differentiation of three zebras sub-species and donkey. Normalized HRM profiles are represented as percent fluorescence for (a) *COI* (b) *cyt b*, and (c) *16S* rRNA markers.

**Figure 5:**
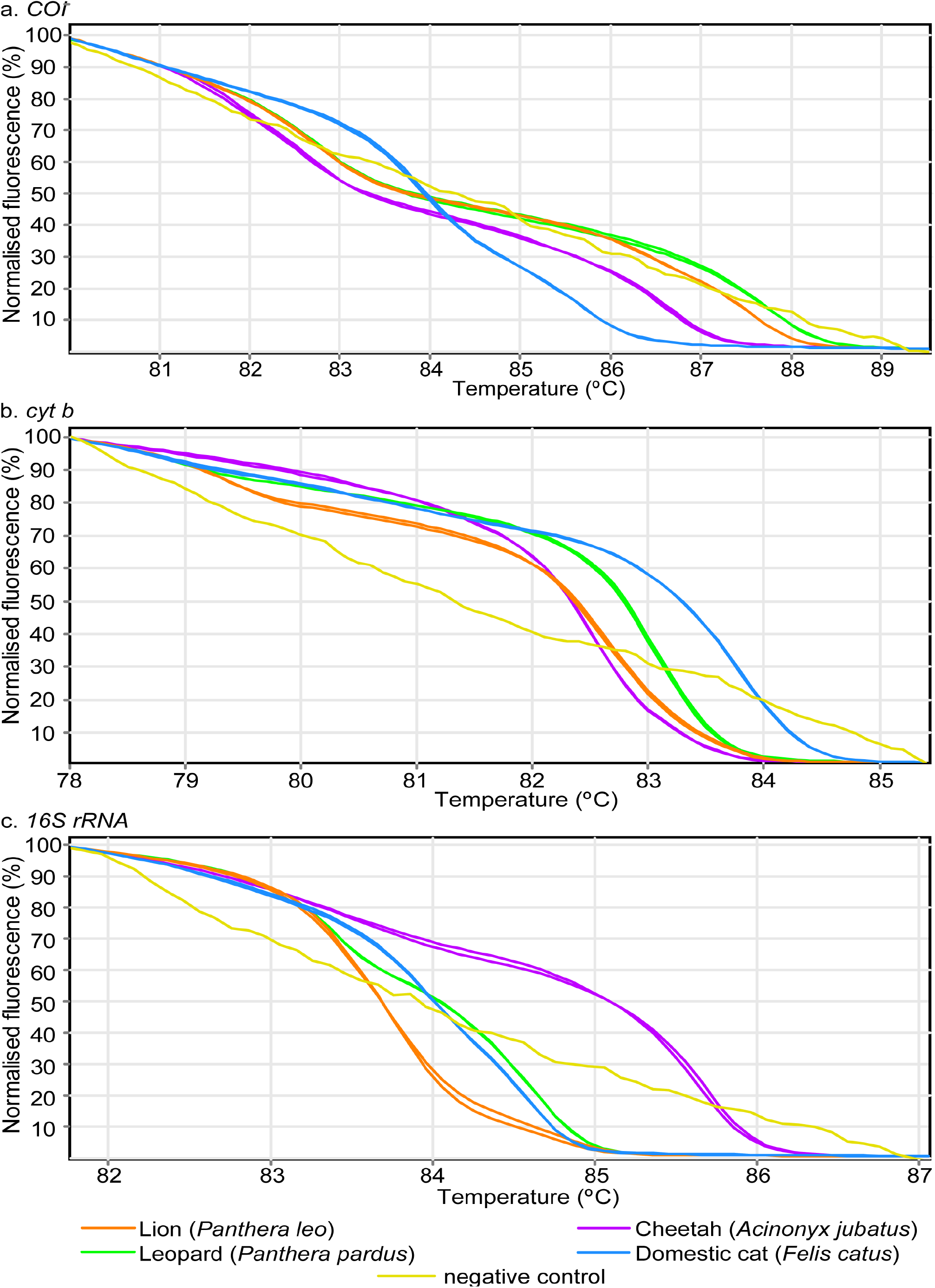
Distinct PCR-HRM profiles for Felidae family species. Normalized HRM profiles are represented as percent fluorescence for (a) *COI* (b) *cyt b*, and (c) *16S* rRNA markers.

**Figure 6:**
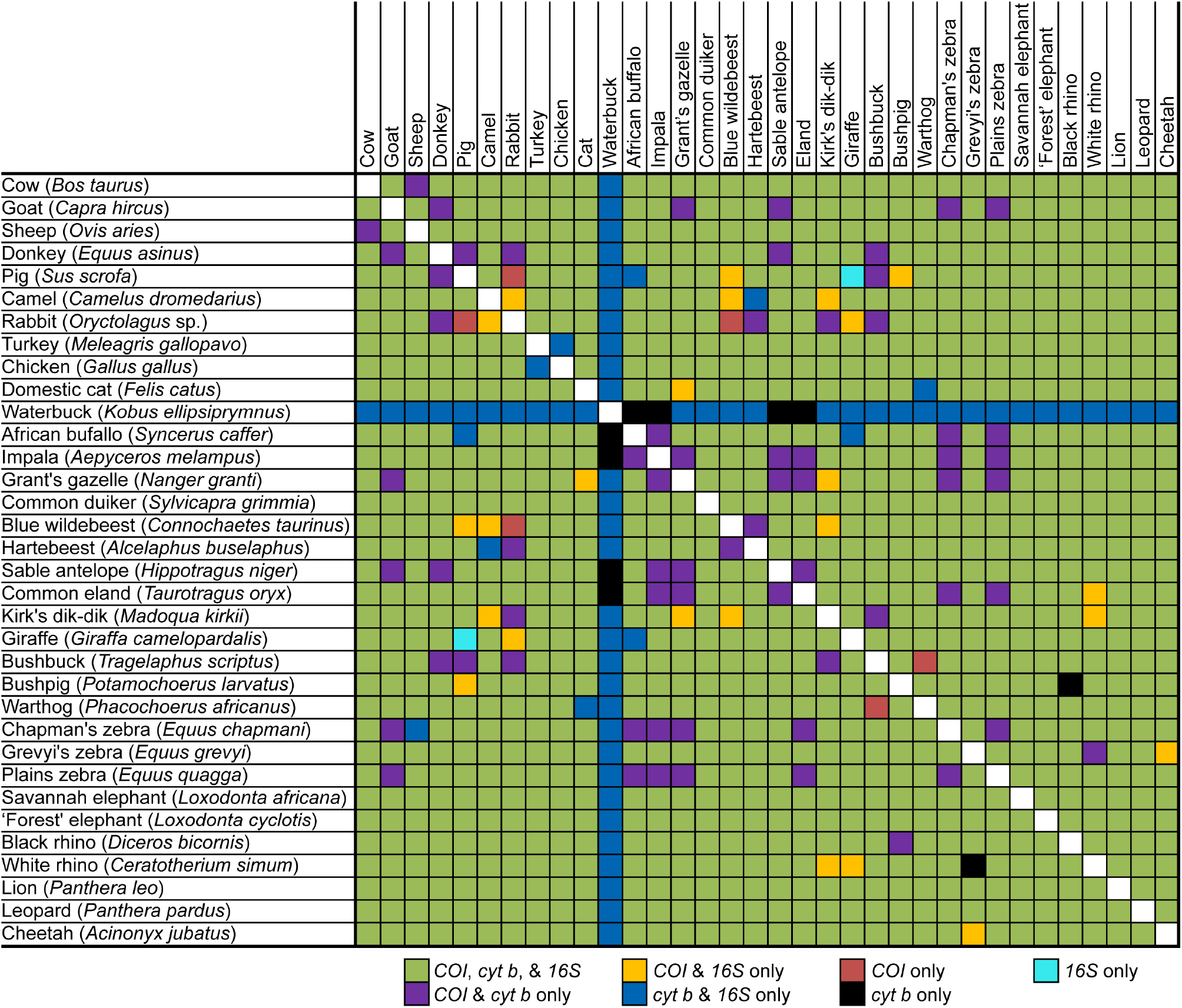
Summary matrix of pair-wise discriminations by PCR-HRM of 34 species and DNA marker resolution. Markers that generated distinct HRM profiles for pair-wise species comparisons are indicated by colours according to the legend.

### Marker discrimination comparison

We did not encounter any species among those tested that could not be distinguished from others by the combined analysis of HRM profiles generated by all the three mtDNA markers. To compare species detection power of molecular markers, the resolution of *COI, cyt b*, and *16S rRNA* PCR-HRMs were compared. Based on 561 pair-wise comparisons of species differentiation (Fig. 6); 39 pairs (7%) could not be distinguished by *COI* PCR-HRM, of which 33 pairs were due to non-amplification of a species (waterbuck) during PCR, and 12 (2.3%) and 33 (6.3%) pairs could not be distinguished by *cyt b* and *16 rRNA* PCR-HRM, respectively. Although PCR-HRM analysis of the *COI* marker was consistently best at resolving species for DNA samples that amplified, giving unique melt curve profiles in shape and T_m_., the *cyt b* and *16S rRNA* markers had better PCR efficiency in all cases for any particular sample, observed by the lower C_T_ values and higher fluorescence values in the melt curve plot. The *cyt b* marker resolved species better than the *16S* marker, which had the highest number of species pairs with similar PCR-HRM profiles. While it is expected that longer PCR products would have more than one melt peak due to the tendency to have multiple melting domains, *cyt b* (~383 bp) and 16S rRNA (~200 bp) PCR products tended to have simple single-peaks compared to *COI* PCR products (~205 bp), for which many samples had multiple peaks, generating a greater diversity of unique HRM profiles.

### Comparison of DNA extraction protocol

The commercially available Qiagen DNeasy Blood and Tissue Kit is used widely across molecular biology laboratories. In comparing this kit with a cheaper laboratory-optimized SDS-proteinase K protocol, we mostly observed similar HRM curve profiles, however with T_m_ shifts ranging from − 0.02 to + 0.4°C, +0.15 to +0.25°C and +0.13 to +0.35°C for *COI, cyt b* and *16S rRNA* markers, respectively, when using the laboratory-optimized protocol compared to the Qiagen protocol (Supp. Fig. 2).

### HRM identification of covertly sampled meat from rural Kenyan butcheries

We covertly sampled 90 meat samples with support of the KWS from butcheries in the Naivasha region of Kenya (0°43’ 0.01” N 36° 26’ 9.28” E, about 77 km from the capital Nairobi). Naivasha is in the vicinity of wild animal conservancies and game parks. We identified one sample That was sold as legal domestic meat (from Kambi Samaki area) to be giraffe bushmeat by PCR-HRM and subsequent COI barcode sequencing confirmation (GenBank accession xxx). The remaining 89 samples consisted of 49 (54.4%) sheep, 29 (32.2%) cattle, eight (8.9%) goats, and two (2.2%) pigs, while one (1.1%) sample failed to amplify. Out of the 17 random samples whose species identity were given by the butcher at the point of sale, six samples (35.3%) sold as goat meat were confirmed by PCR-HRM analysis to be sheep meat (GenBank accession xxx).

## Conclusions

This study clearly demonstrates the utility of PCR-HRM analysis of three mtDNA markers for efficiently differentiating and identifying the vertebrate species origin of unknown tissue samples. Using PCR-HRM, we were able to differentiate domestic and wild animal species native to East Africa by HRM analysis of short *COI*, *cyt b*, and *16S* rRNA gene PCR amplicons. Further, we used the approach to blindly identify illegal giraffe bushmeat among meat samples purchased from rural butcheries, using forensic barcode sequencing only for confirmation purposes. Therefore, the PCR-HRM approach presented here, represents a valuable addition to molecular forensic pipelines for the surveillance of illegal wildlife products, as it eliminates the need for mass barcode sequencing of suspect specimens, most of which tend to be legally traded domestic animal samples. These assays can also be effectively used for biodiversity surveys from the blood-meals of hematophagous invertebrates^28^ and for consumer protection purposes to ensure that meat products for consumption are labeled properly.

We analyzed the PCR-HRM profiles generated by the three DNA markers from 10 domestic and 24 wildlife species endemic to East Africa, demonstrating the capacity of this approach to differentiate a large range of species. While we were able to differentiate all species when considering the combined analysis of HRM profiles generated by all three mtDNA markers, the HRM profiles of individual markers could not be used to distinguish particular vertebrate species pairs, due to the varied resolution strengths of the three markers^29,30^. Though some studies have only considered one^31^ or two^32^ different DNA markers, this study demonstrates increased robustness of species identification by PCR-HRM when using combined analysis of three mtDNA markers. While the reliability of sequencing, as widely applied in species identification, is undoubted, the high costs associated with it may not be sustainable in some instances, especially where large sample sizes are analyzed^17^. PCR coupled to HRM offers a quicker, cheaper, and relatively easy-to-work-with real-time PCR alternative^25^.

By enabling rapid differentiation of commonly consumed domestic species from wildlife vertebrate species, PCR-HRM limits the need for the expensive and time-consuming process of long COI forensic barcode sequencing in surveillance routines, by excluding commonly sold and used domestic species samples from further analysis required to generate forensic evidence for prosecution. This allows for efficient monitoring of potential illegal bushmeat and other wildlife trade products as a more sustainable long-term activity to deter poaching.

We demonstrated the applicability of PCR-HRM analysis to illegal bushmeat surveillance with a small-scale covert surveillance exercise conducted in collaboration with the KWS. Using the three PCR-HRM assays to screen 90 meat samples sold as domestic livestock meat in Naivasha, Kenya, we identified giraffe bushmeat. This was surprising as we expected poaching of much smaller, easier to trap, ruminants to occur more frequently among illegal bushmeat^27^. Poached giraffe meat products are of particular concern as giraffe populations have been declining in the region. The latest update of the International Union of Conservation of Nature (IUCN), *Red List2018-2*, recently added two of the nine sub-species of giraffes to the “Critically Endangered” category. Five out of seven assessed giraffe sub-species are categorised between “Near Threatened” to “Critically Endangered” (IUCN *Red List2018-2*). Further, the sale of illegal bushmeat as livestock meat presents a public health concern as unsuspecting consumers may be exposed to a higher risk of contracting zoonotic diseases^4,5^ by unknowingly consuming bushmeat. Additionally, we identified sheep meat that was sold by local butcheries as goat meat among the covertly sampled meat. This further demonstrates the potential utility of PCR-HRM for surveillance by consumer protection agencies, such as the Kenya Bureau of Standards (KEBS), which in turn, could inform policy formulation and law enforcement.

During the validation of several elephant reference samples, we identified two sets of distinct HRM profiles among the KWS stock samples of Kenyan savannah elephants. We determined, through *COI* barcode sequencing, that some samples amplified mtDNA sequences associated with forest elephant populations, which are thought not to exist in the region^33^. The forest elephant mitochondria amplicons had distinct HRM profiles using all three markers from samples with savannah elephant mitochondria. This finding could represent an artefact of past hybridization between female forest elephants and male savannah elephants^34^, or male forest elephants and female savannah elephants, inferring paternal mtDNA transfer, as suggested in humans by new research^35^ after which the forest elephant mtDNA persisted in East African savannah elephant populations. Therefore, our method can also be used to identify mtDNA variants within populations. Our findings suggest that further screening of savannah elephant samples by PCR-HRM could determine the frequency of forest elephant mtDNA in savannah elephant populations and potentially *vice versa*.

The ability to distinguish by PCR-HRM between diverse ungulates, including buffalo, cow, waterbuck, sheep, goat, and three different zebra species, suggests that the three-marker PCR-HRM method can differentiate even closely related species. We also note that the *cyt b* and *16S rRNA* HRM profiles for cow, goat, sheep, pig, and chicken samples obtained in this study are comparable to those previously obtained from mosquito blood-meals analyses^21^ to determine host feeding preferences, despite the difference in tissue type and PCR cycling conditions. These observations support the overall reproducibility of the method. Nonetheless, we observed melt temperature frame shift generally resulting in slightly higher T_m_ across species when DNA samples were extracted using a laboratory-optimized SDS-proteinase K DNA extraction procedure rather than the commercial Qiagen DNeasy Blood and Tissue Kit, likely due to differences in salt concentrations within the DNA elutes of the different extraction techniques^36^. Therefore, for comparable and reproducible HRM results, all samples should be processed under the same conditions. However, with more affordable HRM-capable thermocyclers entering the market, such as the MIC-4 (Bio Molecular Systems, Australia) and Chai’s Open qPCR (Chai, CA, USA) thermocyclers, cross-platform comparisons may be aided by novel algorithms to harmonize HRM analysis across platforms. By developing online database approaches for automated HRM curve identification across different platforms, data from laboratories can be shared, compared, and classified using machine learning algorithms that may also reduce the need for positive controls within laboratories^37^.

The differences exhibited by the mtDNA markers in their ability to differentiate any two species by PCR-HRM, strongly support the complementarity in using the combination of the three optimized mtDNA markers, which addresses marker-specific shortcomings in differentiating certain vertebrate species. Previous studies also highlighted the importance of marker complementarity in screening mosquitoes for blood-meal sources using HRM^28^. Even though this means that two to three PCR assays must be run to confidently identify a species, the overall time and cost is still cheaper, as the runs can be done simultaneously. Moreover, there was still no need for large-scale sequencing of all PCR amplicons. Only a few representatives with unique peaks need to be selected for species confirmation through DNA sequencing.

The use of short gene targets of 130 bp, which work best with HRM, for barcode identification of species has been shown to be almost as effective as longer barcode sequencing targets^38^ and are more suitable for environmental samples. Nonetheless, consistent with a previously identified marginal positive correlation between amplicon length and species resolution based on *COI* sequences and sequence amplicons of >200 bp^38^, we found that the ¬200 bp *16S rRNA* amplicon had lower HRM resolving power than the *COI* (~205 bp) and *cyt b* (~383 bp) amplicons. The *16S* rRNA amplicon region might have been too short to incorporate sufficient sequence variations required to distinguish species as effectively as the other two markers. The observation that analysis of *COI* and *cyt b* HRM profiles discriminated vertebrate species better than *16S* rRNA HRM profiles is consistent with a study that found *16S* rRNA sequences to be 2.5 times less variable than *COI* and *cyt b* sequences among rodents within the Praomyini tribe^39^. In contrast to previous studies that have investigated the use of HRM analysis to differentiate vertebrate species using different primers to target discrete taxonomic groups^23^, our study used three different sets of universal primers to “globally” differentiate a large repertoire of species. This suggests that its applicability could be much broader than previously published assays.

The power and utility of three-marker HRM analysis is evident and was validated in our double-blind analysis of bushmeat covertly sampled from local butcheries. Out of the 90 samples drawn from the surveillance exercise, we only had to sequence eight representative DNA samples with unique HRM profiles to confirm species identifications. This translates into a 91% reduction in sequencing costs compared to direct sequencing of all the 90 samples. Among the eight samples sequenced, one was identified as giraffe by HRMA and only needed sequencing confirmation for depositing in the GenBank nucleotide database and future forensic prosecution purposes. The other seven were representative sequences of samples with HRM profiles matching those of domestic livestock species. Despite the challenges of sampling during times not favoring the concealed nature of illegal bushmeat trade (from late morning to early evening) and having to deal with mitigating the alerting-appearance of the KWS covert operations team and vehicle, we managed to find one bushmeat specimen among the samples collected. This shows that the problem of illegal bushmeat trade indeed exists in the sampled area.

## Methods

### Samples for optimization

Samples including muscle, blood, and hide were provided by the Kenya Wildlife Service (KWS) forensic laboratory. Some of the domestic species were purchased from local supermarkets or butcheries. We targeted, among others, commonly hunted bushmeat species, and common domestic species (cattle, goat, sheep, donkey, pig, camel, rabbit, turkey, chicken, and cat). KWS meat samples exhibited varying levels of integrity depending on their state at the point of confiscation. After confiscation or sampling, meat and blood had been stored in −40°C freezers in the KWS forensic laboratory. Hide had been stored at room temperature. Samples were grouped mainly according to their taxonomic families, including Bovidae, Equidae, Felidae, Hominidae, Elephantidae, Rhinocerotidae, and Suidae. The numbers of species within each taxonomic family included in the study was limited by the availability of samples (Supp. Table 1).

### Proof-of-concept surveillance of potential bushmeat

Open butcheries were covertly sampled in 22 demarcations within Naivasha (0°43’ 0.01” N 36° 26’ 9.28” E), between 10 am and 4 pm. They included Kambi Daraja, Gilgil, Kinamba, Kasarani, Kihoto, Kamere, Kwa Muya, Kongoni, DCK, Ndabib, Duro, Kabati, Kanjoo Estate, Mirema, Sanctuary, Karagita, Langalanga, Kikopey, Kambi Somali, Kongasis, Mutaita, and Hell’s Gate. Longonot (S00⁰55.096’ E036⁰31.407’) and Mai Mahiu (S00⁰58.966’ E036⁰35.103’) were sampled along the way, between Naivasha and Nairobi. Samples were separately packed and stored for sub-sampling later in the day. We sub-sampled each sample in triplicate into 1.8 ml cryovials, using a sterile scalpel and fresh gloves for every sample. The samples were stored in shippers with liquid nitrogen until extraction.

### Genomic DNA extraction from meat and blood tissue

We extracted genomic DNA from meat and blood using Qiagen’s DNeasy Blood and Tissue Kit (Qiagen, Hannover, Germany) according to the manufacturer’s instructions, with minimal modifications as follows; 25 μl of proteinase K was used for 2-hour incubations of muscle and hard tissue or 1-hour incubations of blood samples. For comparison purposes, we also extracted total DNA from meat and blood using a laboratory optimised protocol, with minor variations based on sample types (meat, blood). Briefly, in 1.5 ml microcentrifuge tubes, about 3 mm^3^ of meat was added to 450 μl of lysis buffer (10 mM Tris (pH 8.1), 0.5% SDS, 5 mM EDTA and 200 μg/ml proteinase K) or 70 μl of blood was topped up to 450 μl with the lysis buffer. Meat samples were then incubated at 65°C in a water bath for 2 hours, whereas blood samples were incubated for one hour. This was followed by the addition of 150 μl of protein precipitation solution (8M Ammonium acetate, 1mM EDTA) at room temperature, then incubation in ice for 15 minutes. The resulting precipitates were centrifuged for 15 minutes at 25000 relative centrifugal force (rcf) in a 5417R Eppendorf centrifuge at 4°C (Eppendorf, Hamburg, Germany). The resultant supernatants were equilibrated to room temperature and then carefully drawn and added into fresh 1.5ml tubes containing 400ml of isopropanol; the pellets were discarded. The mixtures were inverted 100 times followed by centrifugation at room temperature for 15 minutes at 25000 rcf. The resulting supernatants were discarded and 800μl of 70% cold ethanol was added to each pellet, then inverted several times before centrifugation at 25000 rcf for 15 minutes at 4°C. Subsequently, excess ethanol was carefully decanted off and then pellets were inverted over paper towels to air-dry. DNA was resuspended in PCR grade water and stored at −20°C until use. Replicate extractions were performed on samples of species for which available verified specimens were limited to one (see Supp. Table 1).

### Polymerase chain reaction with high-resolution melting analysis

We used primers targeting vertebrate *cyt b*, *16S* rRNA, and *COI* gene fragments (Table 1) in separate single-plex PCRs. Ten microliter PCRs contained 2 μl of pre-formulated 5X HOT FIREPol^®^ EvaGreen^®^ HRM Mix, no ROX (Solis BioDyne, Tartu, Estonia), forward and reverse primers (Macrogen, Europe) at a final reaction concentration of 0.5 μM and 2 μl of DNA template. PCRs were performed in an HRM-capable RotorGene Q thermocycler (Qiagen, Hilden, Germany). Every run had a set of known positive control samples as well as a no-template negative control. The amplification cycling conditions included an initial hold at 95°C for 15 minutes, then 40-45 cycles of denaturation at 95°C for 20 seconds, annealing at 56°C for 20 seconds and extension at 72°C for 30 seconds, followed by a final extension at 72°C for 5 minutes. Immediately after amplification, amplicons were gradually melted at 0.1°C increments from 75°C to 95°C, with fluorescence acquisition every 2 seconds. From these data, graphs of fluorescence against temperature (°C) were generated, which were normalized between zero and 100 fluorescence in the analysis. Every melting profile was observed against the profiles of a set of determined controls, on a per-target basis. Besides the observation of the melting temperatures, the curve shape was also used to differentiate species. PCR-HRM products for *cyt b* and *16S* rRNA were directly purified using ExoSAP-IT (USB Corporation, Cleveland, OH), according to manufacturer’s protocol and sent them for Sanger sequencing at Macrogen, Europe. Since the PCR-HRM target for *COI* lie within the longer barcoding *COI* region they were not sequenced, rather the products from conventional PCR were sequenced.

**Table 1.**
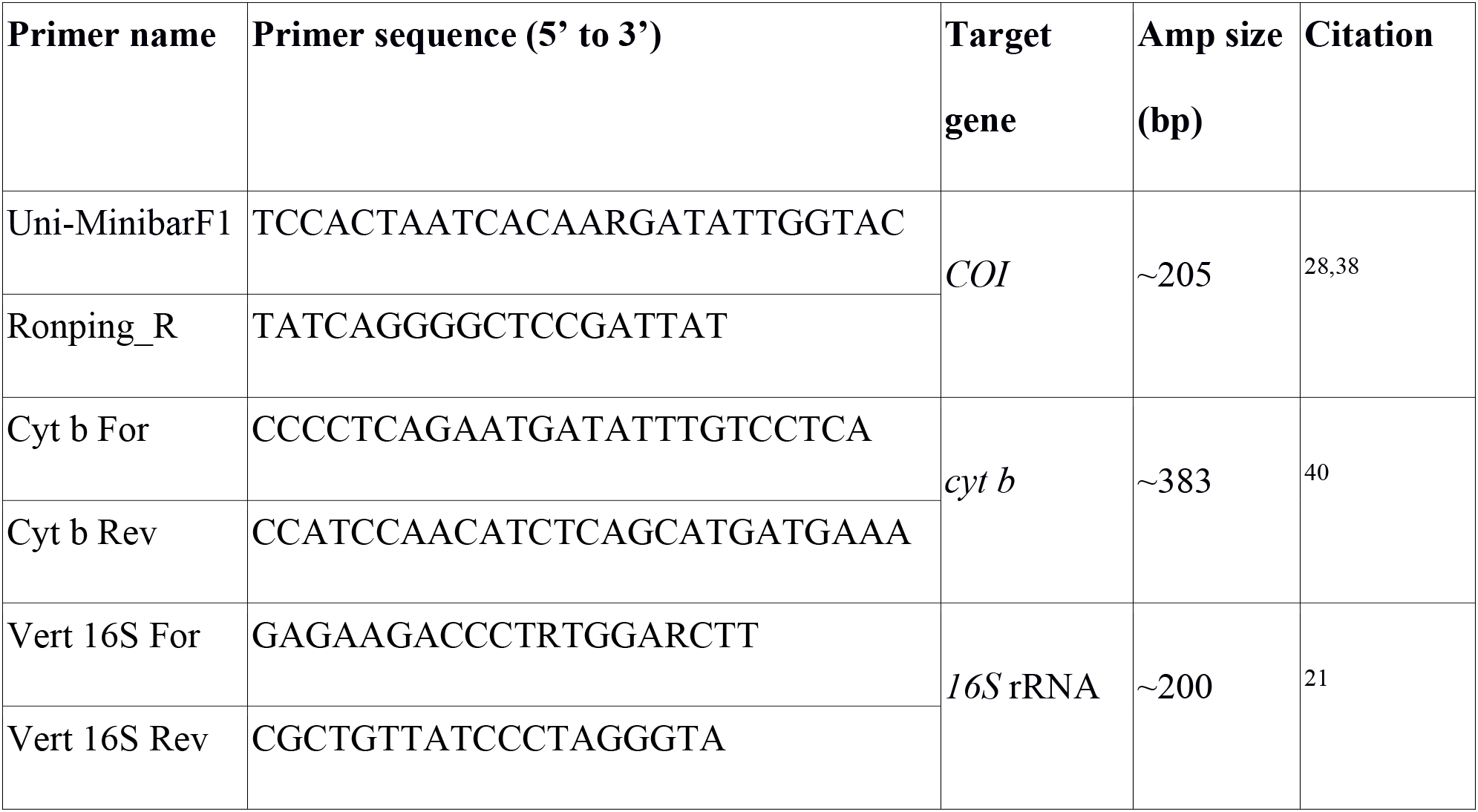
Primers used.

### Long COI polymerase chain reaction

We used conventional PCR for long COI barcode sequencing in 15-μl reaction volumes including 3 μl of 5X HOT FIREPol^®^ Blend Master Mix (Solis BioDyne, Tartu, Estonia), forward and reverse primers at 0.5 μM final concentrations, and 2 μl of template. We used previously described primers VF1d (TCTCAACCAACCACAARGAYATYGG) and VR1d (TAGACTTCTGGGTGGCCRAARAAYCA)^41^ for vertebrate barcoding based on the long (750 bp) COI gene. Each primer was tagged with an M13 tail, TGTAAAACGACGGCCAGT and CAGGAAACAGCTATGAC, forward and reverse respectively, as adapters for sequencing. Amplification conditions included an initial hold at 95°C for 15 minutes, then 40 cycles of denaturation at 95°C for 20 seconds, annealing at 57°C for 30 seconds and extension at 72°C for 60 seconds, followed by a final extension at 72°C for 7 minutes. We electrophoresed PCR products for 45 minutes at 100 V in 2% agarose gels in 1X TAE buffer to ensure proper amplification of target sequences. We then purified amplicons with clear bands using ExoSAP-IT (USB Corporation, Cleveland, OH), according to manufacturer protocol and sent them for Sanger sequencing at Macrogen, Europe.

### Sequence analysis

All sequences were trimmed, edited and analyzed using Geneious v8.1.4 (available from http://www.geneious.com) software^42^ created by Biomatters and queried in GenBank using default BLAST^43^ parameters and aligned sequences obtained with appropriate GenBank reference sequences.

### Blind validation

To validate the three-gene amplicon HRM analysis, we analyzed 90 unknown covert samples through PCR-HRM. From the resultant HRM profiles, we identified unknown samples by comparing their melting profile similarities (melting temperature and curve shape) to the profiles of already known controls. We further confirmed of the identities obtained by HRM analysis through sequencing of the long *COI* fragment and sequence alignment with GenBank reference sequences using Geneious software.

## Acknowledgements

We acknowledge Antoinette Miyunga, (KWS) for help in organizing reference samples and Moses Odhiambo and team (of KWS Naivasha sub office) for their help with the covert surveillance exercise. This study was funded by the United States Agency for International Development Partnerships for Enhanced Engagement in Research (USAID-PEER) cycle 4 awarded to LW, under the USAID grant No. AID-OAA-A-11-00012 sub-awarded by the American National Academy of Sciences (NAS) under agreement No. 2000006204. Additional support was obtained from *icipe* institutional funding from the UK’s Department for International Development (DFID), the Swedish International Development Cooperation Agency (SIDA), the Swiss Agency for Development and Cooperation (SDC), and the Kenyan Government. The funders had no role in the design, data collection, interpretation or decision to submit this publication.

## Author contributions

JV and MYO conceived the study. MYO, MJ, JB, DS, SM, LW, and JV designed the study. DOO, MYO, JWO, and LW coordinated and conducted the covert surveillance activities. DOO performed laboratory experiments. DOO and JV analyzed data and wrote the first draft. All authors contributed to the manuscript editing and approved the final manuscript.

## Supplementary information

**Supplementary Figure 1:**
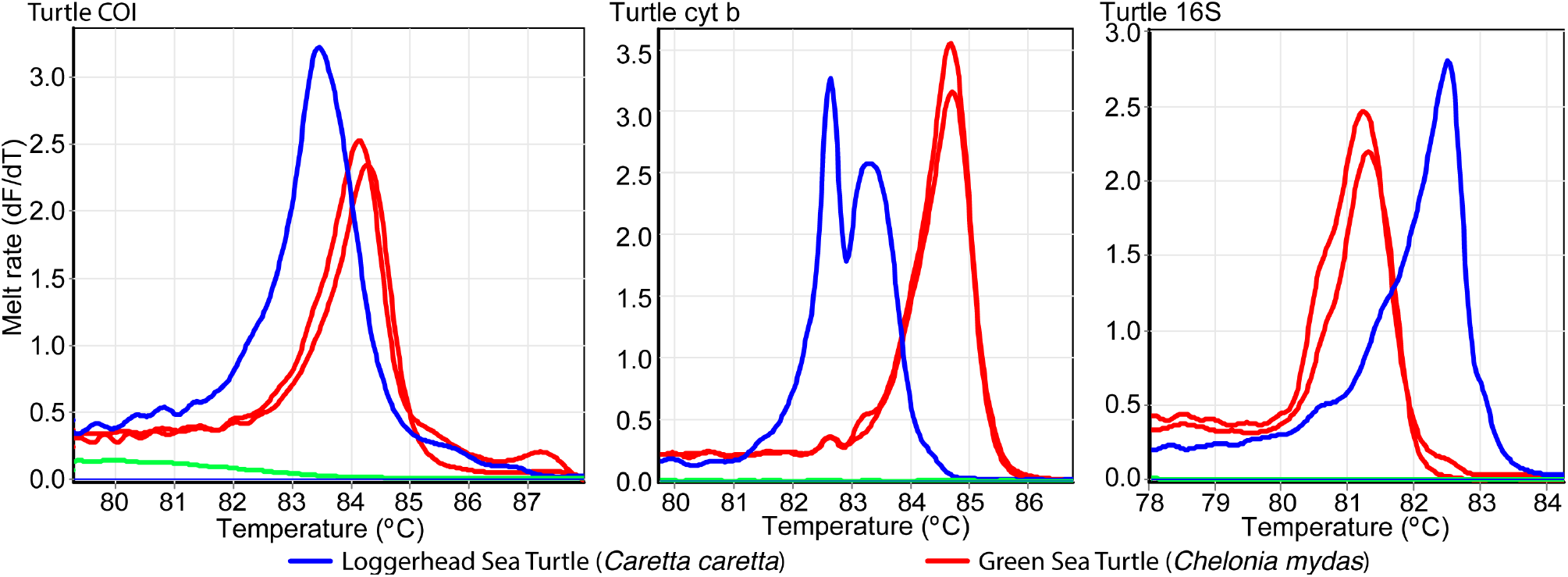
Distinct PCR-HRM melt profiles for Cheloniidae family species. Melt rate profiles are represented as change in fluorescence units with increasing temperatures (dF/dT) for (a) *COI*, (b) *cyt b*, and (c) *16S* rRNA markers.

**Supplementary Figure 2:**
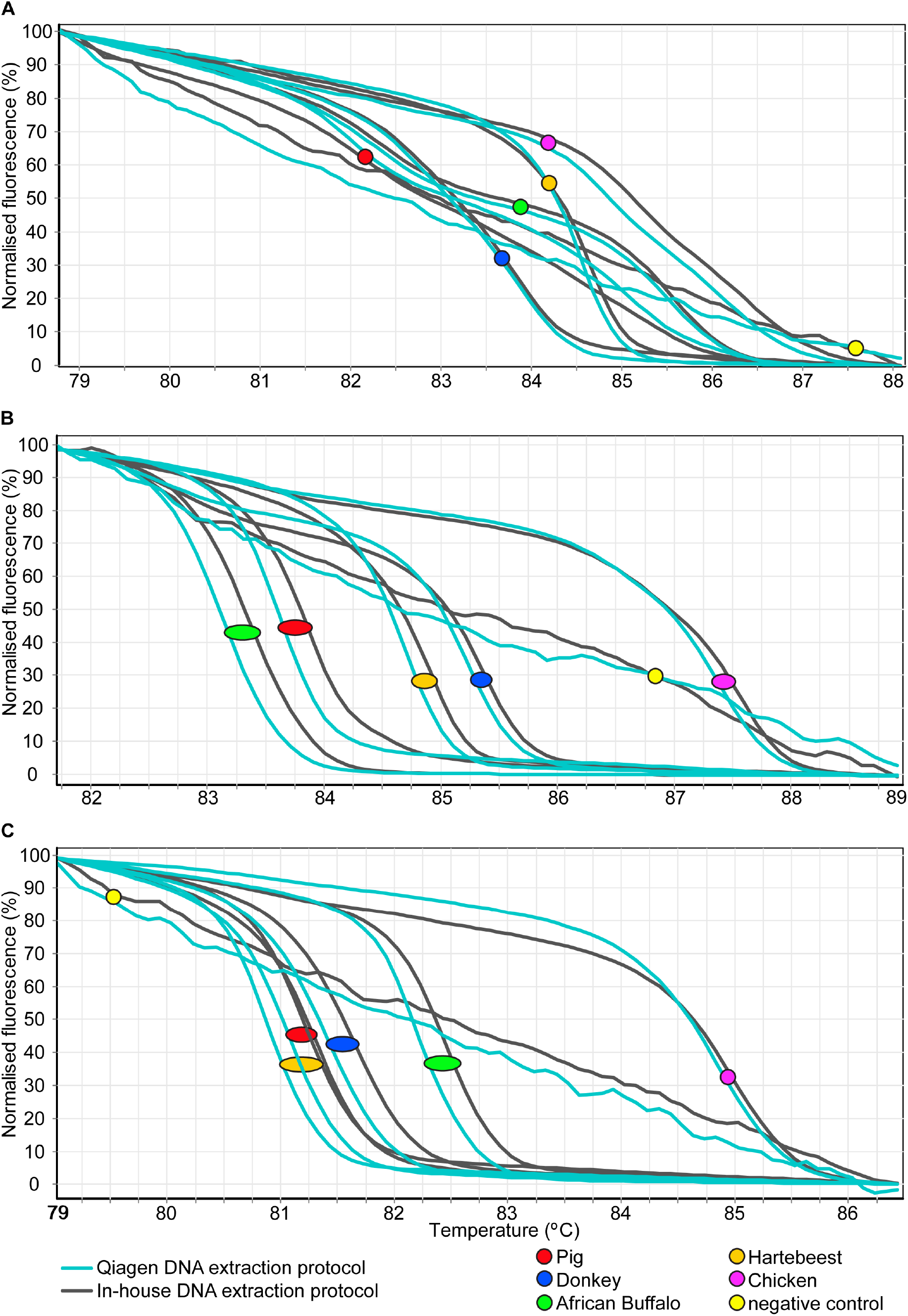
Normalized HRM profiles of representative reference samples extracted using different protocols. Normalized HRM profiles are represented as percent fluorescence for (a) *COI* (b) *cyt b* and (c) *16S* rRNA markers.

**Supplementary Table 1.** Samples used in the HRM reference optimizations.

| **Family** | **Species** | **Common name** | **Tissue** | **Blood** | **Total** |
| --- | --- | --- | --- | --- | --- |
| Bovidae | *Bos taurus* | Cow | 2 | 4 | 6 |
| Bovidae | *Capra hircus* | Goat | 1 | 1 | 2 |
| Bovidae | *Ovis aries* | Sheep | 1 | 1 | 2 |
| Equidae | *Equus asinus* | Donkey | 1 | 0 | 1 |
| Suidae | *Sus scrofa* | Pig | 1 | 0 | 1 |
| Camelidae | *Camelus dromedarius* | Camel | 0 | 1 | 1 |
| Leporidae | *Oryctolagus cuniculus* | Rabbit | 1 | 0 | 1 |
| Phasianidae | *Meleagris gallopavo* | Turkey | 1 | 0 | 1 |
| Phasianidae | *Gallus gallus* | Chicken | 1 | 1 | 2 |
| Felidae | *Felis catus* | Domestic cat | 1 | 0 | 1 |
| Bovidae | *Kobus ellipsiprymnus* | Waterbuck | 3 | 0 | 3 |
| Bovidae | *Syncerus caffer* | African buffalo | 4 | 4 | 8 |
| Bovidae | *Aepyceros melampus* | Impala | 4 | 0 | 4 |
| Bovidae | *Nanger granti* | Grant's gazelle | 1 | 0 | 1 |
| Bovidae | *Sylvicapra grimmia* | Common duiker | 3 | 0 | 3 |
| Bovidae | *Connochaetes taurinus* | Blue wildebeest | 5 | 2 | 7 |
| Bovidae | *Alcelaphus buselaphus* | Hartebeest | 1 | 1 | 2 |
| Bovidae | *Hippotragus niger* | Sable antelope | 0 | 2 | 2 |
| Bovidae | *Tragelaphus oryx* | Eland | 2 | 0 | 2 |
| Bovidae | *Madoqua Kirkii* | Kirk's dik-dik | 1 | 0 | 1 |
| Bovidae | *Tragelaphus scriptus* | Bushbuck | 1 | 2 | 3 |
| Giraffidae | *Giraffa camelopardalis* | Giraffe | 4 | 2 | 6 |
| Suidae | *Potamochoerus porcus* | Bushpig | 2 | 0 | 2 |
| Suidae | *Phacochoerus africanus* | Warthog | 2 | 2 | 4 |
| Equidae | *Equus chapmani* | Chapman's zebra | 2 | 0 | 2 |
| Equidae | *Equus grevyi* | Grevy's zebra | 0 | 2 | 2 |
| Equidae | *Equus quagga* | Plain zebra | 2 | 2 | 4 |
| Elephantidae | *Loxodonta africana* | Savannah elephant | 2 | 4 | 6 |
| Elephantidae | *Loxodonta cyclotis* | 'Forest' elephant | 0 | 2 | 2 |
| Rhinocerotidae | *Diceros bicornis* | Black rhino | 0 | 4 | 4 |
| Rhinocerotidae | *Ceratotherium simus* | White rhino | 0 | 5 | 5 |
| Felidae | *Panthera leo* | Lion | 3 | 3 | 6 |
| Felidae | *Panthera pardus* | Leopard | 0 | 2 | 2 |
| Felidae | *Acinonyx jubatus* | Cheetah | 0 | 3 | 3 |
| Cheloniidae | *Chelonia mydas* | Green sea turtle | 4 | 0 | 4 |
| Cheloniidae | *Caretta caretta* | Logger head sea turtle | 1 | 0 | 1 |
| **Total:** |  |  | 57 | 50 | 107 |

